# Deciphering the orthorhombic crystal structure of a novel NEIL1 nanobody with pseudo-merohedral twinning

**DOI:** 10.1101/2023.08.07.552313

**Authors:** Marlo K. Thompson, Nidhi Sharma, Aishwarya Prakash

## Abstract

Nanobodies or VHHs (Variable Heavy domains of Heavy chain) are single domain antibodies that comprise three antigenic complementary determining regions (CDR). Nanobodies are used in numerous scientific applications including, bio-imaging, diagnosis, therapeutics, and macromolecular crystallography. We obtained crystals of a ∼14 kDa nanobody specific for the NEIL1 DNA glycosylase (hereafter called A5) in 0.5 M ammonium sulfate, 0.1 M sodium citrate tribasic dihydrate pH 5.6, and 1.0 M lithium sulfate monohydrate from the Crystal HT Hampton Research screen that were further optimized. Here, we describe the structure determination and refinement of the A5 crystals to a resolution of 2.1 Å. The data collected were complicated by the presence of anisotropy and twinning, and while initial space group determination pointed to a higher apparent tetragonal crystal system, the data statistics suggested twinning, placing the crystal in an orthorhombic system. Twinning was confirmed by the Padilla and Yeates test, H-test, and Britton test based on local intensity differences with a twin fraction of 0.4. Molecular replacement produced the best solution in the orthorhombic space group P2_1_2_1_2 with four molecules in the asymmetric unit and we were able to model over 96% of the residues in the electron density with a final R_work_ and R_free_ of 0.1988 and 0.2289 upon refinement.

**Synopsis:** The crystal structure of a specific nanobody against NEIL1 was determined to 2.1 Å. The structure was ultimately solved in an orthorhombic space group after diffraction data analysis revealed mild anisotropy as well as pseudo-merohedral twinning

## 1. Introduction

Despite an increase in structures solved by cryo-electron microscopy (Cryo-EM), X-ray crystallography remains one of the most utilized methods to determine the three-dimensional structures of macromolecules such as proteins^1,2^, accounting for nearly 75% of the structures released since 2020 (rcsb.org, visited Jan 30^th^, 2023)^3^. In many cases, however, the crystallization of a native full-length protein is not a straightforward process as many proteins harbor disordered regions that are often not conducive to the crystallization process^4^. A common way to surmount this hurdle is to obtain crystals by truncating the disordered regions of protein in order to obtain a crystallizable fragment with the caveat that valuable information contained within these regions is lost. To overcome this issue, there has been an increase in the use of stabilizing partners to co-crystallize with a native protein containing a degree of disorder in order to alter the crystal lattice formation in a thermodynamically favored manner^4^. One such chaperone is the single variable domain from camelid heavy-chain only antibodies, better known as VHH or nanobodies^5^. Specific examples exist in the literature where nanobodies have been successfully used to aid in the crystallization and structure determination of their target proteins^6-8^.

In contrast to the conventional heterotetrameric antibody, nanobodies have many advantages that make them appealing for use in a variety of applications from structural to *in cellulo* studies. Nanobodies can be purified in large volumes in a standard laboratory using recombinant expression in a bacterial host. Furthermore, the nanobody sequences can be easily tailored for specific uses such as bio-imaging, therapeutics, diagnostics, and crystallography (reviewed in Muyldermans, S., 2021)^9^. Nanobodies have been reported to have higher stability than fragment antigen-binding regions (Fabs) or single-chain fragments (scFvs) and can bind targets with nanomolar to micromolar affinity (K_D_)^10,11^.

We developed nanobodies against the NEIL1 DNA glycosylase, a DNA repair enzyme that participates in the oxidized lesion recognition and excision steps of the DNA base excision repair pathway. The full-length mature NEIL1 enzyme comprises 390 amino acids where over a quarter of the protein (∼100 residues at the C-terminus) is disordered and participates in various protein-protein interactions involving replication and repair factors. Crystal structures of unliganded NEIL1 as well as NEIL1 bound to lesion-containing DNA substrates have been obtained, but they lack the C-terminal disordered region^12,13^. Furthermore, the field of NEIL1 biology suffers owing to low endogenous cellular NEIL1 expression levels as well a lack of reliable commercially available antibodies against the protein. While co-crystals of the larger NEIL1-nanobody complex have yet to be obtained, in this current work, we obtained diffraction-quality crystals of the anti-NEIL1 nanobody (also referred to as A5) after extensive optimization of the protein sequence, affinity tags used during purification, and the crystallization conditions. We identified pseudo-merohedral twinning within the data where the crystals belong to an orthorhombic crystal system and the twin-law supports higher metric symmetry. This type of twinning was observed previously with crystals of a variable domain of SAG173-04^14^, and the crystals of the enzyme, Dicer^14,15^.

## 2. Materials and methods

### 2.1. Identification of Anti-NEIL1 nanobodies

Nanobodies specific for NEIL1 were identified using the Hybrigenics Services SAS (available in the public domain at www.hybrigenics-services.com, Paris, France) synthetic human nanobody library by Yeast 2-Hybrid methods^16^. The NEIL1 enzyme was used as a bait protein in the LexA and Gal4 screens, as described previously^17^. The LexA and Gal4 screens surveyed 98 and 69 million clones and returned 3 and 33 positive clones, respectively. Clones corresponding to 10 unique nanobody sequences were delivered by Hybrigenics. The sequence of the A5 nanobody was identified in both screens with multiple clones and was used for all subsequent work described herein.

### 2.2. Plasmids, Nanobody Expression, and Purification

Nanobody sequences with C-terminal Flag and 10x histidine (His) tags were cloned into a pEBY vector. To optimize expression yields and crystallization, gene blocks containing A5 fused with alternative purification tags (Genscript; Table 1) were inserted via standard cloning techniques and digested with *Nco*I and *Not*I (NEB) restriction enzymes. The A5 constructs were transformed into Rosetta2 (DE3) pLysS *Escherichia coli* cells (Novagen) and selected on 100 µg/ml ampicillin and 100 µg/ml chloramphenicol plates. A single colony was used to generate a primary culture with overnight growth in 100 ml terrific broth supplemented with 0.4% glycerol, 1% glucose, 100 µg/ml ampicillin, and 100 µg/ml chloramphenicol, shaking at 37 °C. A larger secondary culture was generated using 20 ml of the primary culture in 1 L terrific broth supplemented with 0.4% glycerol, 0.1% glucose, 100 µg/ml ampicillin, and 100 µg/ml chloramphenicol, shaking at 37 °C. Protein expression was induced with 0.5 mM isopropyl-D-thiogalactopyranoside (IPTG) overnight at 28 °C. Cells were pelleted by centrifugation at 14,000 x g for 10 min.

**Table 1:** Protein constructs of A5 with various tags that were purified and subjected to crystallizations trials are displayed. His-tags, Flag-tags, TEV cleavage sites, and residues used as linkers or cloning sites are colored as green, blue, red, pink respectively.

| Construct | Vector | Complete amino-acid sequence of the construct produced | Result |
| --- | --- | --- | --- |
| A5-10His-Flag | pEBY | MAEVQLQASGGGFVQPGGSLRLSCAASGATSERDIMGWFRQAPGKE<br>REFVSAISYEPGGWHYYADSVKGRFTISRDN SKNTVYLQMNSLRA<br>EDTATYYCAWQEIMYGEAYWRYWGQGTQVTVSSAAALEAHHHHHH<br>HHHHGAADYKDDDDK | Small, thin crystal clusters |
| 6His-tev-A5 | pEBY | MHHHHHHENLYFQ SMAEVQLQASGGGFVQPGGSLRLSCAASGATSE<br>RDIMGWFRQAPGKEREFVSAISYEPGGWHYYADSVKGRFTISRDN SKNTVYLQMNSLRAEDTATYYCAWQEIMYGEAYWRYWGQGTQVTVSS | Insoluble |
| 6His-A5 | pEBY | MHHHHHHSMAEVQLQASGGGFVQPGGSLRLSCAASGATSERDIMGW<br>FRQAPGKEREFVSAISYEPGGWHYYADSVKGRFTISRDN SKNTVYLQMNSLRAEDTATYYCAWQEIMYGEAYWRYWGQGTQVTVSS | Insoluble |
| A5-6His | pEBY | MAEVQLQASGGGFVQPGGSLRLSCAASGATSERDIMGWFRQAPGKE<br>REFVSAISYEPGGWHYYADSVKGRFTISRDN SKNTVYLQMNSLRA<br>EDTATYYCAWQEIMYGEAYWRYWGQGTQVTVSSGHHHHHH | Small, thin crystal clusters |
| A5-tev-6His | pEBY | MAEVQLQASGGGFVQPGGSLRLSCAASGATSERDIMGWFRQAPGKE<br>REFVSAISYEPGGWHYYADSVKGRFTISRDN SKNTVYLQMNSLRA<br>EDTATYYCAWQEIMYGEAYWRYWGQGTQVTVSSENLYFQSHHHHHH | Individual rod crystals |

The cell pellet was resuspended in a buffer containing 50 mM HEPES pH 7.4, 500 mM NaCl, 1 mM ethylenediaminetetraacetic acid (EDTA), 1 mM dithiothreitol (DTT), 1 mM phenylmethylsulfonyl fluoride (PMSF), and cOmplete (EDTA-free protease inhibitor cocktail; Sigma), sonicated, and centrifuged at 25,000 x g. Clarified cell lysates were incubated with pre-equilibrated TALON Superflow beads (Cytiva) for 2 hours at 4 °C. Beads were washed with 10 column volumes of wash buffer (50 mM HEPES pH 7.4, 100 mM NaCl, and 1 mM DTT) and resuspended for cleavage of the 6x His-tag by addition of 0.8 mg TEV protease per 15 mL of resuspended resin. The protein elutes in the flow through and is subsequently subjected to size exclusion chromatography using a Superdex 200 (GE Healthcare) in a buffer containing 50 mM HEPES (pH 8.2), 200 mM NaCl, 10% (v/v) glycerol, and 1 mM DTT. Fractions were pooled, concentrated, and flash-frozen aliquots were stored at -80 °C.

### 2.3. Crystallization and data collection

We crystallized A5 with various purification tags (Table 1; and Fig. 1) until we obtained crystals that produced quality diffraction data. Original crystallization drops were prepared by mixing 0.5 µL protein (6.8 mg/mL) and 0.5 µL reservoir solution containing 0.1 M sodium citrate pH 5.6, 1.2 M lithium sulfate, and 0.5 M ammonium sulfate. Trays were incubated at 14 °C and crystals were obtained in 3 days. A crystal was macro-seeded into a new drop made using 0.5 µL protein (6.875 mg/mL), 0.4 µL of the previously described reservoir solution, and 0.1 µL of beaded seeds from a similar condition (0.1 M sodium citrate pH 5.6, 1.4 M lithium sulfate, and 0.125 M ammonium sulfate). The macro-seeded crystal was collected after 7 days and mounted onto a MiTeGen micro-loop, cryoprotected (0.1 M sodium citrate pH 5.6, 1.2 M lithium sulfate, and 0.5 M ammonium sulfate, and 20% glycerol), and flash cooled in liquid nitrogen. Crystals were screened on our home source (Bruker D8 Quest) and a complete dataset was collected on the X-ray Operations and Research beamline 22-ID at the Advanced Photon Source, Argonne National Laboratory.

**Figure 1:**
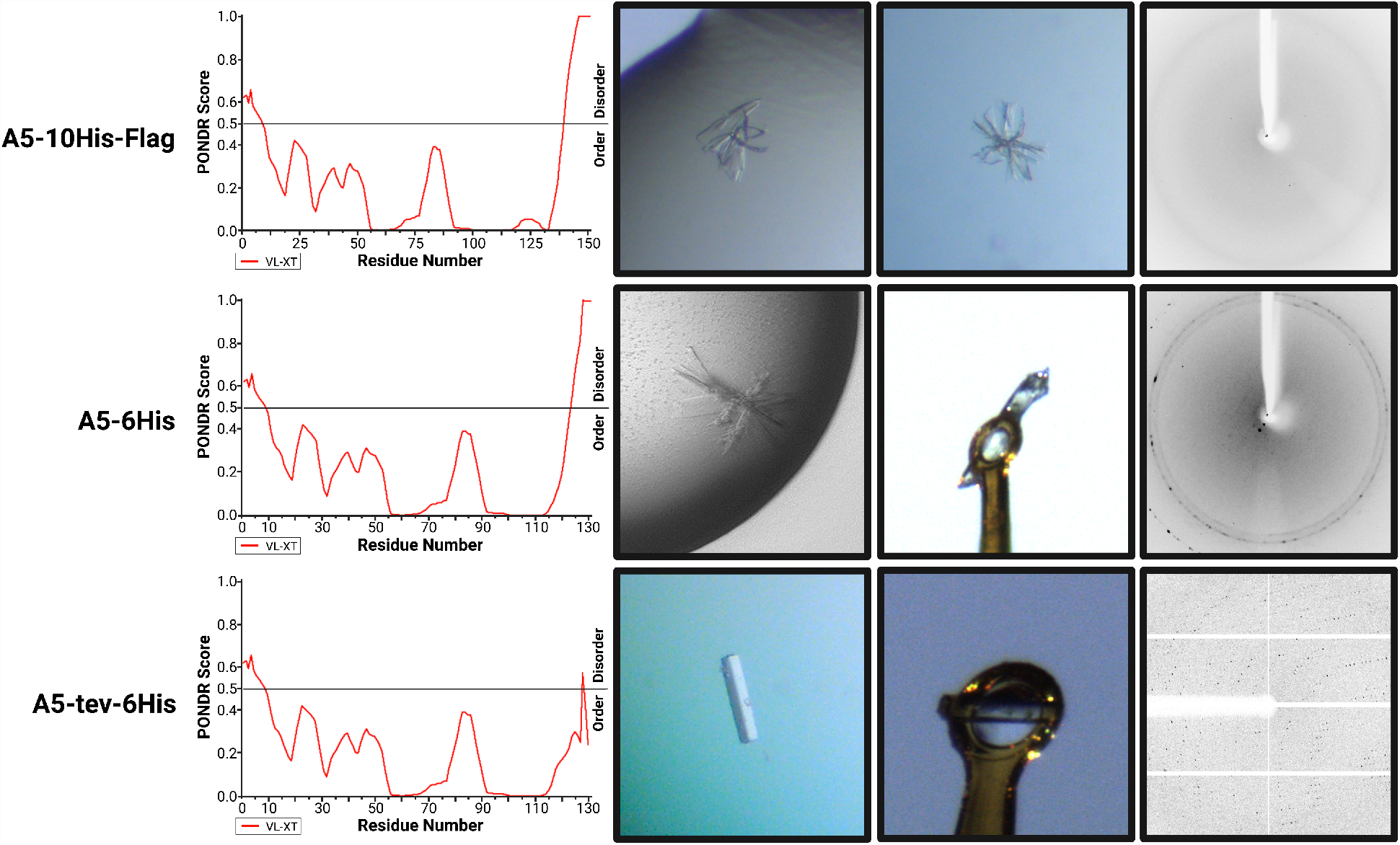
Increased crystal quality with decreasing construct disorder. A5-10His-Flag (top row) PONDR analysis (left) predicted the largest region of disorder at the C-terminus. A5-10His-Flag produced small, clustered crystals that were unable to produce a diffraction pattern (right). PONDR predicted A5-6His (middle row) to have a shorter disordered region at the C-terminus than the 10His-Flag construct above. The A5-6His protein produced slightly larger, plate-like crystal clusters that produced low-resolution diffraction. The cleavable A5-tev-6His protein (bottom row) was predicted to have minimal disorder within the C-terminal region. The cleaved A5-tev-6His produced individual, 3-dimentional rod-shaped, crystals that diffracted to 2.08 Å.

### 2.4. Data analysis and phase determination by molecular replacement

Data were processed using the Proteum suite^18^. Detection of twinning was performed in *phenix.xtriage*^19^. Determination of the twin law governing the pseudo-merohedrally twinned crystal was performed following proper space group determination. Molecular replacement was performed using *phenix.phaser*^20^ using the structural coordinates of the anti-RHOB nanobody (PDB: 6SGE chain B)^15^. The atomic coordinates of the anti-RHOB nanobody were extracted and modified with *phenix.sculptor*^21^ to truncate any sidechains that aligned with non-identical residues in A5 (77% sequence identity). The modified anti-RHOB nanobody coordinates were used as the search model for molecular replacement in *phenix.phaser*.

### 2.5. Model building and refinement

Models were built using *Coot* (version 0.9.8.5) and refined using *phenix.refine*^19^. The twin operator (*-h, l, k*) was applied during each round of refinement. All water and sulfate molecules were added in *Coot*. The quality of the final model was validated by *phenix.molprobity*, with 97.7% of residues in the preferred Ramachandran regions^22^. Structural coordinates were deposited to the protein data bank (PDB) with the code 8FTT. All structural figures were generated with *MacPyMOL* (The PyMOL Molecular Graphics System, Version 1.7.6.2; Schrödinger, LLC).

## 3. Results and Discussion

### 3.1. Optimization of initial crystallization and data processing

We obtained small, clustered, plate-like, fragile crystals that did not produce a diffraction pattern with the original 10His-Flag tagged A5 protein (Table 1, Fig. 1). Further truncation of the 10His-Flag tag to a 6His tag yielded larger clusters of crystals, albeit, with poor diffraction (Fig. 1). We ultimately obtained single rod-shaped crystals with reduced levels of disorder, as predicted by the Predictor of Natural Disordered Regions (PONDR) (Fig. 1)^23^, with a construct comprising a TEV cleavable C-terminal 6His-tag. We observed diffraction to ∼3 Å upon screening using our Bruker D8 Quest home source. A complete dataset was subsequently acquired at the APS beamline 22-ID. Initial close observation of the spot intensities indicated the appearance of smearing caused by partially overlapping reflections (Fig. 2) and was an indication of potential twinning.

**Figure 2:**
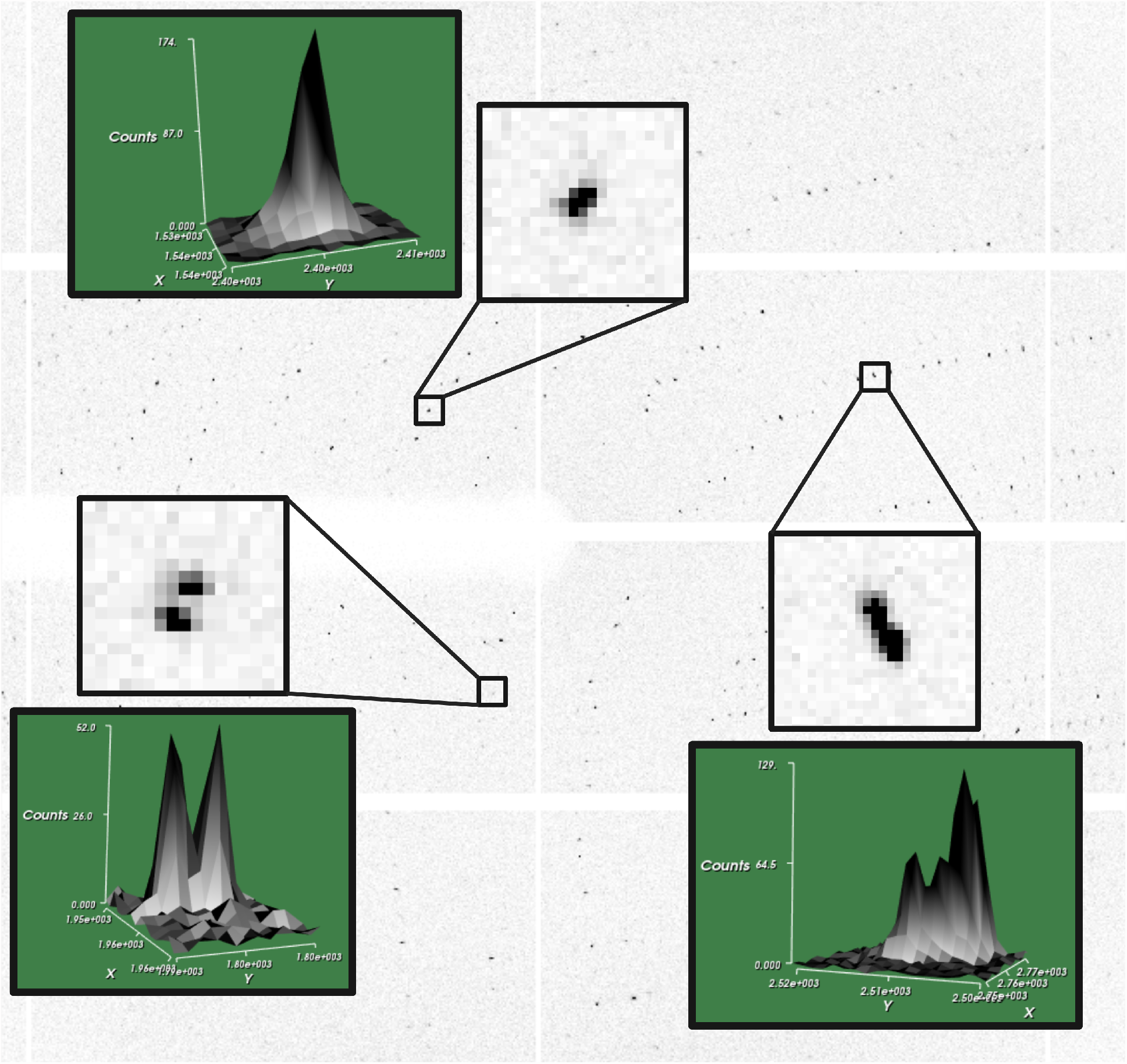
The diffraction pattern of A5 reveals possibility for twinning. The twinning of A5 can be seen in the diffraction pattern produced by the crystals and the observed partially overlapping spots. The enlarged view of the diffraction pattern (insets) display a single spots (top), minimally overlapping spots (bottom left), and split spots (bottom right). The green box adjacent to each spot is the corresponding 3D peak profile, with pixel intensity represented as counts.

### 3.2. Space group determination

In order to analyze the quality of the data, we evaluated the unmerged data using *phenix.xtriage*. Based on initial results, the space group was assigned as P4_2_22. Wilson analysis suggested that the data were mildly anisotropic, which was confirmed by STARANISO^24^. Other diagnostic tests suggested twinning within the data, with a multivariate Z score of 13.470 (Tables 2,3). Results from the Padilla-Yeates algorithm^25^ for twin detection, Britton estimation of the twin fraction^26^, and the H-plot analysis^27^ can be found in Table 2. Furthermore, the placement of our data into the space group P4_2_22, was unexpected as only ∼0.01% of soluble monomeric proteins fall into this space group.^28-30^ Our initial MR attempt did not produce a correct solution. While the statistics looked promising (49≤LLG≤60; 7≤TFZ≤8), the resulting coordinates did not pack satisfactorily without severe clashes between neighboring molecules (Table 3)^31^. With the lack of a *bona fide* molecular replacement solution, and the likelihood of twinning within the data, we examined lower symmetry space group options not supported by the Laue group 4/*mmm*.

**Table 2:**
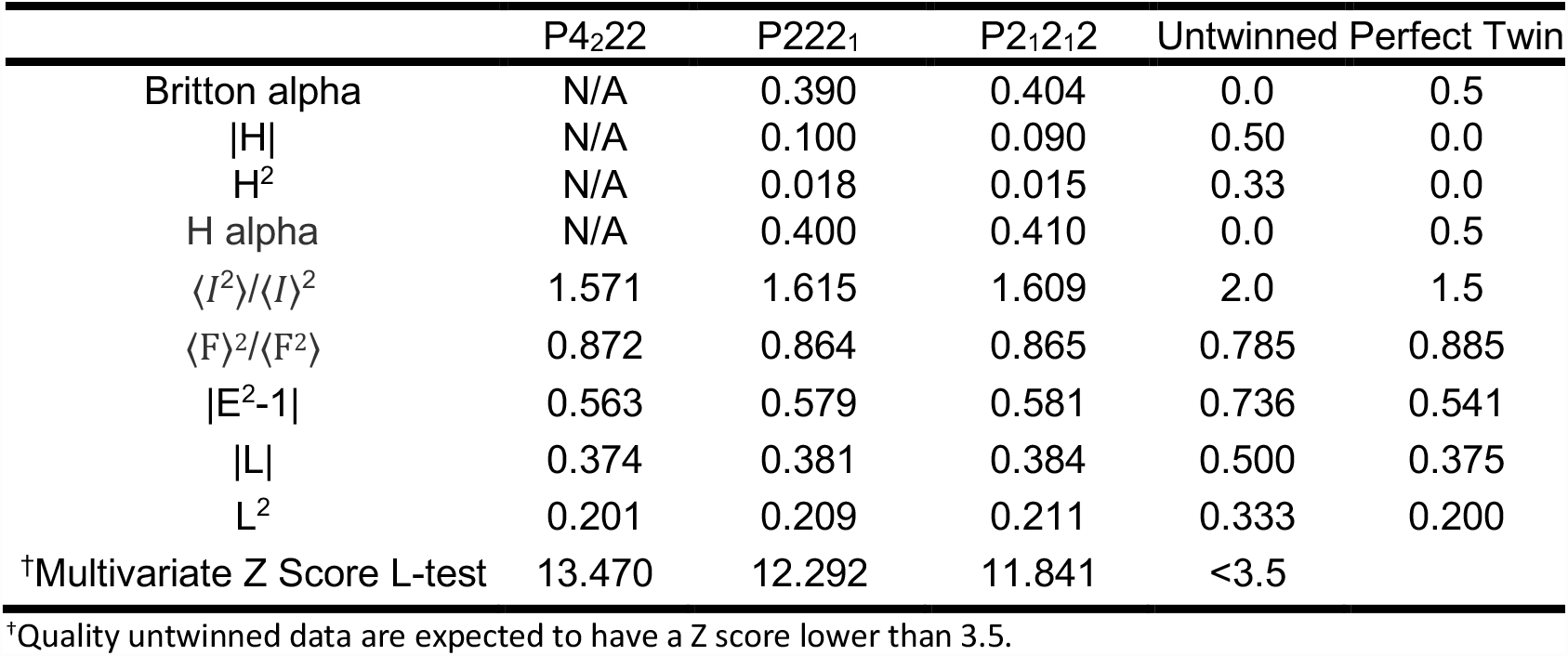
Twinning Statistics Independent of Twin Laws.

**Table 3:** MR statistics from various space groups.

| | $P4_222$ | $P222_1$ | $P2_12_12$ |
| --- | --- | --- | --- |
| Solutions Found | 48 | 1 | 1 |
| Top LLG | 58.760 | 185.035 | 655.133 |
| Top TFZ | 7.3 | 9.6 | 17.9 |

We reprocessed the data using a primitive orthorhombic cell. Interestingly, the merging statistics did not drastically change (Table 4). Analysis indicated that the space group was P222_1_ and that the data were anisotropic with a strong possibility of twinning (Table 2). The Matthew’s coefficient was calculated to be 2.53 Å^3^ Da^-1^ (51.4% solvent), suggesting five A5 molecules per asymmetric unit (ASU). Based on the Matthew’s coefficient, we ran MR searching for five component copies and returned a single solution with an LLG of 185.035 and TFZ of 9.6. However, the MR solution contained only three molecules within the ASU, which corresponds to a Matthews coefficient of 4.21 Å^3^ Da^-1^ (70.8% solvent). Furthermore, we did not observe any direct interactions between neighboring molecules in the ASU. We then broadened our MR search by screening all possible space groups associated with the point group 222 and varying the number of component copies. A single solution with four components was obtained in the space group P22_1_2_1_ with a LLG of 516.491 and TFZ of 20.0. The four components were placed in the cell without any severe clashes or overlaps. The unit cell was transformed and the space group was assigned to the standard setting of P2_1_2_1_2, we reprocessed the data in the space group P2_1_2_1_2 and obtained a final MR solution with a LLG of 655.133 and a TFZ of 17.9 (Table 5).

**Table 4:** Data collection from various space groups.

|  | P4<sub>2</sub>22 | P2<sub>1</sub>2<sub>1</sub>2 |
| --- | --- | --- |
| Resolution (Å) | 38.00-2.10 (2.16-2.10) | 40.27-2.08 (2.14-2.08) |
| Resolution* (Å) |  |  |
| *a* * | 2.2 | 1.9 |
| *b* * | 2.2 | 2.4 |
| *c* * | 2.0 | 2.3 |
| Redundancy | 14.0 (14.9) | 6.9 (7.0) |
| Completeness (%) | 100.0 (99.9) | 99.3 (93.9) |
| Rmerge | 0.153 (1.454) | 0.120 (1.825) |
| Rmeas | 0.159 (1.506) | 0.130 (1.987) |
| *I*/σ(*I*) | 12.0 (2.1) | 10.6 (1.3) |
| Unit-cell parameters |  |  |
| *a* (Å) | 75.9932 | 121.8190 |
| *b* (Å) | 75.9932 | 76.0320 |
| *c* (Å) | 121.855 | 75.7685 |
| Molecules per ASU | 2 | 4 |
| CC1/2 | 0.998 (0.901) | 0.998 (0.678) |
| Wilson *B* (Å<sup>2</sup>) | 33.29 | 34.67 |
| $\langle I^2 \rangle / \langle I \rangle^2$ | 1.571 | 1.609 |

Values in parentheses are for the highest resolution shell. $R_{merge} = \sum ||I - \langle I \rangle| / \sum I$ , where $\langle I \rangle$ is the average intensity from multiple observations of symmetry-related reflections. \* Anisotropy resolution limits calculated by STARANISO are provided along *a*\*, *b*\*, and *c*\* for the datasets are indicated.

**Table 5:** Data collection, phasing, and refinement statistics for A5.

| **A5 (PDB: 8FTT)** |  |
| --- | --- |
| Beamline | Advanced Photon Source |
| Wavelength (Å) | 1.00 |
| Space group | P 21 21 2 |
| Unit-cell parameters (Å, °) | a=121.8190, b=76.0320, c=75.7685<br>α=90, β=90, γ=90 |
| Molecules per asymmetric unit | 4 |
| **Data collection statistics** |  |
| Resolution range (Å) | 40.27-2.08 (2.14-2.08) |
| Unique reflections | 42462 (3094) |
| Redundancy | 6.9 (7.0) |
| Rmerge | 0.120 (1.825) |
| Rmeas | 0.130 (1.968) |
| Rpim | 0.049 (0.724) |
| CC1/2 | 0.998 (0.678) |
| Overall I/σ | 10.6 (1.3) |
| Completeness (%) | 99.30 (93.9) |
| **MR phasing statistics** |  |
| Top LLG | 655.133 |
| Top TFZ | 17.9 |
| **Refinement Statistics** |  |
| Reflections used in refinement | 42436 (4024) |
| Reflections used for R-free | 4280 (420) |
| R<sub>work</sub> (%) | 0.1988 (0.3165) |
| R<sub>free</sub> (%) | 0.2289 (0.3289) |
| CC<sub>work</sub> | 0.888 (0.355) |
| CC<sub>free</sub> | 0.824 (0.305) |
| Number of non-hydrogen atoms | 3995 |
| Macromolecules | 3939 |
| Ligands | 30 |
| Solvent | 26 |
| r.m.s.d values |  |
| Bond length (Å) | 0.003 |
| Bond angles (°) | 0.50 |
| B-factor (Å<sup>2</sup>) |  |
| Wilson B | 42.82 |
| Macromolecules | 42.72 |
| Ligands | 57.73 |
| Water | 40.98 |
| Ramachandran plot |  |
| Ramachandran favored (%) | 97.34 |
| Ramachandran allowed (%) | 2.66 |
| Ramachandran outliers (%) | 0.00 |
| Clashscore | 6.10 |

$R_{merge} = \sum ||I - \langle I \rangle| / \sum I$ , where $\langle I \rangle$ is the average intensity from multiple observations of symmetry-related reflections. $R_{work}$ and $R_{free} = \sum ||F_o| - |F_c|| / \sum ||F_o||$ , where $F_o$ and $F_c$ are the observed and calculated structure factor amplitudes, respectively. $R_{free}$ was calculated with 10% of the reflections not used in refinement. MR, molecular replacement. Values for the highest resolution shell are shown in parentheses.

### 3.3. Twinning and refinement

In an orthorhombic system with twinned domains, the unit-cell must be made up of two axes nearly identical in length, as observed with the current A5 crystals where b≈c (121.8190, 76.0320, 75.7685;), making the lattice approximately tetragonal^14^. The subsequent relationship between the twin domains creates pseudo-4-fold symmetry along a, via the twin operation (-*h, l, k*). The second screw axis is masked by the superposition of the two short axes. Ultimately, using refinement in P2_1_2_1_2 with the twin operator (-*h, l, k*), we obtained a twin fraction of 0.43, and a finalized A5 model with an R_work_ and R_free_ of 0.1988 and 0.2289, respectively (Table 5).

### 3.4. Overall structure of the A5 nanobody

The overall three-dimensional structure of A5 is displayed in Fig. 3A. We were able to model over 96% of the residues into the electron density with no Ramachandran outliers upon refinement. A5 displays a canonical immunoglobulin fold (Ig-fold) with a β-sandwich formed by two antiparallel β-sheets connected by loops, turns, and pi helices (residues 65-70 and 90-94) (Fig. 3A-C). CDRs 1–3 (residues 29-35, 55-61, and 101-112, respectively) form the epitope recognition site, with CDR3 extending the β-strand framework of the core β-sandwich. The A5 core contains a canonical disulfide bond between β-strands 3 and 9, which proceed CDR1 and CDR3, respectively (residues C24 and C99; Fig. 3A-C). The presence of this conserved disulfide bond lends stability to the nanobodies by increasing their thermal denaturation temperatures^32,33^.

Based on our MR solution, we observe four molecules (Chains A-D) within the ASU displayed in Fig. 3D. Each molecule in the ASU interacts with a total of five neighboring molecules, two within the ASU and three symmetry mates, with the exception of Chain A which only contacts two symmetry mates (Fig. 4). Chain A will be used as an example to describe the network of interactions that occur between molecules. CDR3 (residue Y106 and W111) and the following framework (residue W114) of Chain A interacts with CDR2 (residue Y55) of Chain B and the TEV recognition site (residues N126, L127, F129, and Q130) of Chain C (Fig. 4A). Residue W60 in CDR2 of chain A forms a hydrogen bond with Q119 of Chain C (Fig. 4B). The loop containing CDR2 of chain A (residue S54) forms an indirect interaction with chain C (residue Q119) via a water molecule (Fig. 4B). CDR2 (residue E56) and the loop between β7 and β8 (residue N77) of Chain A interacts with CDR3 (residue I104 and Y106) of the symmetry mate 1 (s.m.1) of Chain C (Fig. 4C). β9 (residue T96) and β11 (residues Q119, V120, and T121) of Chain A form direct and indirect interactions with CRD2 (residues E56 and W60) and β5 (residue S54) of a second symmetry mate (s.m. 2) of Chain C (Fig. 4D). The C-terminus of Chain A (residues S123, S124, E125, L127, F129, Q130) interacts with CDR3 of Chain C s.m.2 (residues Q102, A109, W111, and Y113) both directly and indirectly (Fig. 4E). We consistently observe interactions that involve CDR2, CDR3, and the TEV recognition site of each molecule. The amino acids involved in the interactions within each region differ between chains (Fig. 4A, E). However, one interaction that is conserved between chains is the hydrogen bond between the carbonyl of W111 and the amine of F129 of a neighboring chain. Chain A is lacking a bond between the C-terminal residues of neighboring symmetry mates. Q130 of Chain B forms a single hydrogen bond with Q130 of Chain C, while Q130 of Chain D forms a salt bridge with another molecule of Chain D.

**Figure 3:**
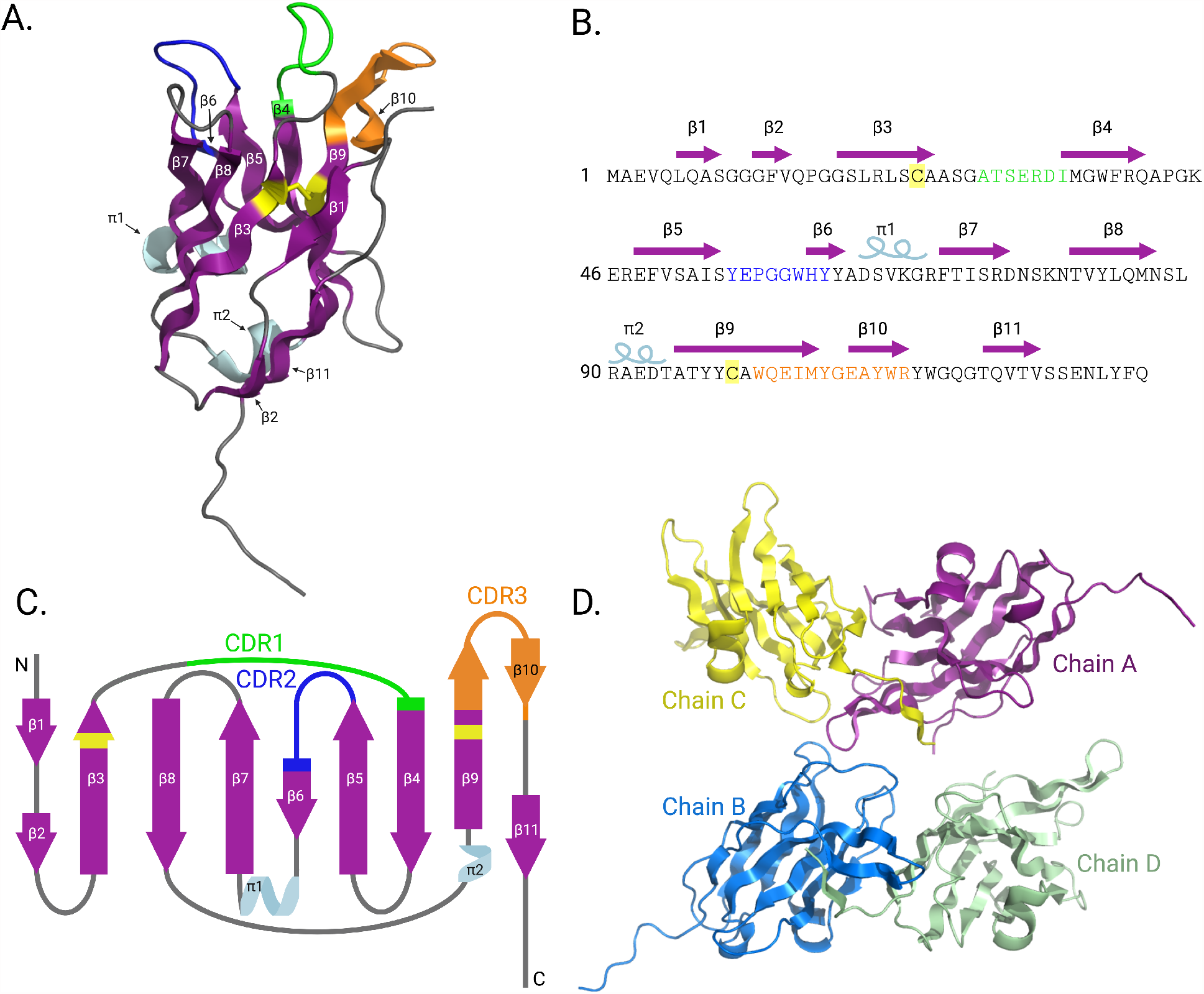
The overall three-dimensional structure of A5. A. The overall three-dimensional structure of A5 colored as follows: β-strands, purple arrows; pi-helices, light blue; CDR1, green; CDR2, blue; CDR3, orange. The β-strands (β1-11) and pi-helices (π1-2) are labeled in sequential order. The disulfide bond formed between residues C24 and C99 is highlighted in yellow. B. The A5 sequence with the major secondary structures displayed by purple arrows (β-strands) and coils (pi-helices). C. Topology map of A5. D. Organization of the four molecules (chains A-D) within the asymmetric unit (ASU).

**Figure 4:**
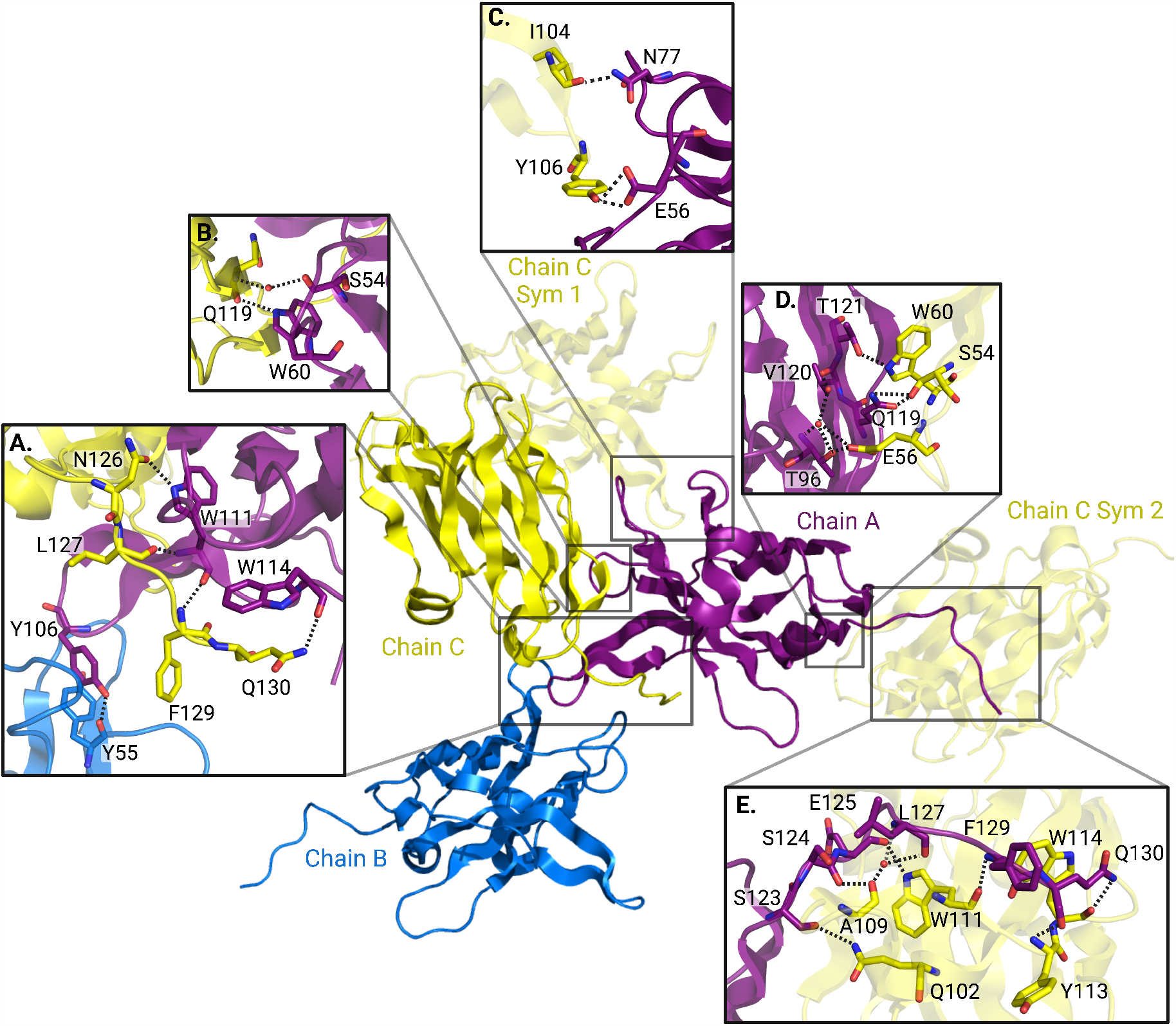
Stabilizing interactions between Chain A and neighboring molecules. A-B. H-bond interaction network at the interface of Chains A, B and C in the asymmetric unit. C-E. H-bond interaction network highlighting interactions between Chain A and symmetry mates of Chain C.

## 4. Conclusions

Nanobodies are beneficial from a structural perspective to bind to regions of proteins that are inaccessible to Fabs, to stabilize dynamic proteins^34-36^, mask flexible regions^37^, stabilize specific protein conformations in multi-domain proteins^38^, and stabilize macromolecular complexes^39^. We will utilize our knowledge regarding A5 to enable structure determination of the full-length NEIL1 complex, as well utilize the nanobodies generated in applications including tracking localization of NEIL1 in tissue culture models. The generation and structural characterization of the NEIL1 nanobodies will revolutionize our understanding of NEIL1 biology.

## Acknowledgements

We would first like to thank Dr. Matthew Benning (Bruker) for being intimately involved with all steps of the project including data processing, steps taken to solve the structure, and providing critical feedback on the manuscript. We would also like to thank Caleb Stratton, a graduate student at University of South Alabama for assisting us with remote data collection at the APS and Dr. Brian Eckenroth (University of Vermont) for reviewing the manuscript and proving critical feedback. Use of the APS was supported by the U.S. Department of Energy, Office of Science, Office of Basic Energy Sciences, under Contract No. DE-AC02-06CH11357. MKT, NS, and AP are supported by a grant from the National Institutes of Environmental Health Sciences (NIEHS), Outstanding New Environmental Scientist (ONES) R01 grant #R01ES030084 to AP. Startup funds provided to AP by the University of South Alabama Health Mitchell Cancer Institute are also acknowledged.

## References Cited

1 Kwan, T. O. C., Axford, D. & Moraes, I. Membrane protein crystallography in the era of modern structural biology. Biochem Soc Trans 48, 2505–2524, doi:10.1042/BST20200066 (2020).

2 Powell, H. R. A beginner’s guide to X-ray data processing. The Biochemist 43, 46–50, doi:10.1042/bio_2021_124 (2021).

3 Berman, H. M. et al. The Protein Data Bank. Nucleic Acids Res 28, 235–242, doi:10.1093/nar/28.1.235 (2000).

4 Koide, S. Engineering of recombinant crystallization chaperones. Curr Opin Struct Biol 19, 449–457, doi:10.1016/j.sbi.2009.04.008 (2009).

5 Hamers-Casterman, C. et al. Naturally occurring antibodies devoid of light chains. Nature 363, 446–448, doi:10.1038/363446a0 (1993).

6 Duhoo, Y. et al. Camelid nanobodies used as crystallization chaperones for different constructs of PorM, a component of the type IX secretion system from Porphyromonas gingivalis. Acta Crystallogr F Struct Biol Commun 73, 286–293, doi:10.1107/S2053230X17005969 (2017).

7 Kumar, H. et al. Crystal Structure of a ligand-bound LacY-Nanobody Complex. Proc Natl Acad Sci U S A 115, 8769–8774, doi:10.1073/pnas.1801774115 (2018).

8 Hansen, S. B., Laursen, N. S., Andersen, G. R. & Andersen, K. R. Introducing site-specific cysteines into nanobodies for mercury labelling allows de novo phasing of their crystal structures. Acta Crystallogr D Struct Biol 73, 804–813, doi:10.1107/S2059798317013171 (2017).

9 Muyldermans, S. Applications of Nanobodies. Annu Rev Anim Biosci 9, 401–421, doi:10.1146/annurev-animal-021419-083831 (2021).

10 Pardon, E. et al. A general protocol for the generation of Nanobodies for structural biology. Nat Protoc 9, 674–693, doi:10.1038/nprot.2014.039 (2014).

11 Chabrol, E. et al. VHH characterization.Recombinant VHHs: Production, characterization and affinity. Anal Biochem 589, 113491, doi:10.1016/j.ab.2019.113491 (2020).

12 Doublie, S., Bandaru, V., Bond, J. P. & Wallace, S. S. The crystal structure of human endonuclease VIII-like 1 (NEIL1) reveals a zincless finger motif required for glycosylase activity. Proc Natl Acad Sci U S A 101, 10284–10289, doi:10.1073/pnas.0402051101 (2004).

13 Zhu, C. et al. Tautomerization-dependent recognition and excision of oxidation damage in base-excision DNA repair. Proc Natl Acad Sci U S A 113, 7792–7797, doi:10.1073/pnas.1604591113 (2016).

14 Brooks, C. L. et al. Pseudo-symmetry and twinning in crystals of homologous antibody Fv fragments. Acta Crystallogr D Biol Crystallogr 64, 1250–1258, doi:10.1107/S0907444908033453 (2008).

15 MacRae, I. J. & Doudna, J. A. An unusual case of pseudo-merohedral twinning in orthorhombic crystals of Dicer. Acta Crystallogr D Biol Crystallogr 63, 993–999, doi:10.1107/S0907444907036128 (2007).

16 Moutel, S. et al. NaLi-H1: A universal synthetic library of humanized nanobodies providing highly functional antibodies and intrabodies. Elife 5, pdoi:10.7554/eLife.16228 (2016).

17 Jang, E. R. et al. HUWE1 is a molecular link controlling RAF-1 activity supported by the Shoc2 scaffold. Mol Cell Biol 34, 3579–3593, doi:10.1128/MCB.00811-14 (2014).

18 PROTEUM3 v. 2019.11-0 (Bruker, 2019).

19 Adams, P. D. et al. PHENIX: building new software for automated crystallographic structure determination. Acta Crystallogr D Biol Crystallogr 58, 1948–1954, doi:10.1107/s0907444902016657 (2002).

20 Storoni, L. C., McCoy, A. J. & Read, R. J. Likelihood-enhanced fast rotation functions. Acta Crystallogr D Biol Crystallogr 60, 432–438, doi:10.1107/S0907444903028956 (2004).

21 Bunkoczi, G. & Read, R. J. Improvement of molecular-replacement models with Sculptor. Acta Crystallogr D Biol Crystallogr 67, 303–312, doi:10.1107/S0907444910051218 (2011).

22 Chen, V. B. et al. MolProbity: all-atom structure validation for macromolecular crystallography. Acta Crystallogr D Biol Crystallogr 66, 12–21, doi:10.1107/S0907444909042073 (2010).

23 Xue, B., Dunbrack, R. L., Williams, R. W., Dunker, A. K. & Uversky, V. N. PONDR-FIT: a meta-predictor of intrinsically disordered amino acids. Biochim Biophys Acta 1804, 996–1010, doi:10.1016/j.bbapap.2010.01.011 (2010).

24 Tickle, I. et al. STARANISO Global Phasing Ltd. Cambridge, UK (2018).

25 Padilla, J. E. & Yeates, T. O. A statistic for local intensity differences: robustness to anisotropy and pseudo-centering and utility for detecting twinning. Acta Crystallogr D Biol Crystallogr 59, 1124–1130, doi:10.1107/s0907444903007947 (2003).

26 Britton, D. Estimation of twinning parameter for twins with exactly superimposed reciprocal lattices. Acta Crystallographica Section A 28, 296–297 (1972).

27 Yeates, T. O. Simple statistics for intensity data from twinned specimens. Acta Crystallogr A 44 (Pt 2), 142–144, doi:10.1107/s0108767387009632 (1988).

28 Parsons, S. Introduction to twinning. Acta Crystallogr D Biol Crystallogr 59, 1995–2003, doi:10.1107/s0907444903017657 (2003).

29 Chruszcz, M. et al. Analysis of solvent content and oligomeric states in protein crystals--does symmetry matter? Protein Sci 17, 623–632, doi:10.1110/ps.073360508 (2008).

30 Gaur, R. K. Frequency distribution of space groups in soluble and membrane proteins and their complexes. Acta Crystallogr F Struct Biol Commun 77, 187–191, doi:10.1107/S2053230X21005719 (2021).

31 McCoy, A. J., Read, R. J., Bunkóczi, G. & Oeffner, R. D. PhaserWiki: Molecular Replacement, < https://www.phaser.cimr.cam.ac.uk/index.php/Molecular_Replacement> (2009).

32 Liu, H., Schittny, V. & Nash, M. A. Removal of a Conserved Disulfide Bond Does Not Compromise Mechanical Stability of a VHH Antibody Complex. Nano Lett 19, 5524–5529, doi:10.1021/acs.nanolett.9b02062 (2019).

33 Hagihara, Y., Mine, S. & Uegaki, K. Stabilization of an immunoglobulin fold domain by an engineered disulfide bond at the buried hydrophobic region. J Biol Chem 282, 36489–36495, doi:10.1074/jbc.M707078200 (2007).

34 De Bruyn, P. et al. Nanobody-aided crystallization of the transcription regulator PaaR2 from Escherichia coli O157:H7. Acta Crystallogr F Struct Biol Commun 77, 374–384, doi:10.1107/S2053230X21009006 (2021).

35 Korotkov, K. V., Pardon, E., Steyaert, J. & Hol, W. G. Crystal structure of the N-terminal domain of the secretin GspD from ETEC determined with the assistance of a nanobody. Structure 17, 255–265, doi:10.1016/j.str.2008.11.011 (2009).

36 Lam, A. Y., Pardon, E., Korotkov, K. V., Hol, W. G. J. & Steyaert, J. Nanobody-aided structure determination of the EpsI:EpsJ pseudopilin heterodimer from Vibrio vulnificus. J Struct Biol 166, 8–15, doi:10.1016/j.jsb.2008.11.008 (2009).

37 Loris, R. et al. Crystal structure of the intrinsically flexible addiction antidote MazE. J Biol Chem 278, 28252–28257, doi:10.1074/jbc.M302336200 (2003).

38 Rasmussen, S. G. et al. Structure of a nanobody-stabilized active state of the beta(2) adrenoceptor. Nature 469, 175–180, doi:10.1038/nature09648 (2011).

39 Rasmussen, S. G. et al. Crystal structure of the beta2 adrenergic receptor-Gs protein complex. Nature 477, 549–555, doi:10.1038/nature10361 (2011).

